# Approximate Bayesian Computation for Inferring Waddington Landscapes from Single Cell Data

**DOI:** 10.1101/2023.09.03.556134

**Authors:** Yujing Liu, Stephen Y. Zhang, Istvan T. Kleijn, Michael P.H. Stumpf

## Abstract

Single cell technologies allow us to gain insights into cellular processes at unprecedented resolution. In stem cell and developmental biology snapshot data allows us to characterise how the transcriptional state of cells changes between successive cell types. Here we show how approximate Bayesian computation (ABC) can be employed to calibrate mathematical models against single cell data. In our simulation study we demonstrate the pivotal role of the adequate choice of distance measures appropriate for single cell data. We show that for good distance measures, notably optimal transport distances, we can infer parameters for mathematical models from simulated single cell data. We show that the ABC posteriors can be used to characterise parameter sensitivity and identify dependencies between different parameters, and to infer representations of the Waddington or epigenetic landscape, which forms a popular and interpretable representation of the developmental dynamics. In summary, these results pave the way for fitting mechanistic models of stem cell differentiation to single cell data.

## 1 Introduction

While developmental biology has largely progressed through observational studies, from the beginning of the 20th century this seemingly intricate, bewilderingly complex, yet robust, process has also fascinated mathematicians. Of particular importance has been C.H. Waddington’s epigenetic landscape[24]. Originally intended as a metaphor for developmental processes, the idea of a landscape, has continued to guide thinking among biologists and mathematicians; the Fields medalist René Thom, for example, was interested in developmental biology as a potential manifestation of catastrophe theory.

Over the past decade there has been a resurgence in interest from mathematicians, developmental biologists, and bioengineers in the epigenetic or Waddington landscape. We can use it as a mathematical and computational tool to reason about and even predict cell fates[31, 52]. Recent studies [12, 40, 36, 16] have given us qualitative insights into the fundamental dynamics underlying differentiation at the cellular level. In a deterministic framework it was shown that even fundamental developmental dynamics can be understood in terms of elegant mathematical models of small gene regulation networks [36]. But to account for the (experimentally quantifiable) randomness prevailing among sub-cellular molecular processes we must extend such analyses to incorporate stochastic processes. This has become especially important because stochastic effects can change the dynamics not just quantitatively but qualitatively, profoundly reshaping the manifold on which dynamics occur. In specific, when external multiplicative noise are present, cell states represented by attractors in the landscape can be destroyed or created [23, 16].

While the mathematical analysis of dynamical systems has achieved a high level of maturity, our ability to challenge mathematical models of developmental systems with data is lagging behind both the mathematical theory and our capability to probe developmental systems in experiment. Technological advances in single cell biology provide us with exquisitely detailed snapshots of the transcriptomic states of single cells [30]; and before long we will also be able to collect single cell protein data of the required quality and quantity [37]. And while descriptive and statistical analyses of single cell data have progressed in lock-step with experimental technologies and new data, mechanistic modelling in light of data has progressed more slowly.

Landscapes for dynamical system models of developmental systems are typically formulated by a specific manner through thermodynamic approaches with detailed chemical assumptions about kinetics of gene transcription factors [43]. But the inference of parameters underpinning the landscapes remains an open problem. It is well known that variation of the parameters of a dynamical system can lead to changes in the cellular state or even the structure of the landscapes: for example, at some critical value of parameters, creation or destruction of attracting states may occur which leads to different landscape structures. Such changes in dynamical system due to the variation of parameters are known as bifurcations [25, 12] and are a central phenomenon of study in the dynamical systems literature. Therefore, to reconstruct the landscapes that agree experimental observations, it is necessary to have parameters adequately inferred.

Coupling single cell data to modelling has been challenging as it has been difficult to e.g. estimate reaction rate parameters for dynamical systems models. Here we explore the use of approximate Bayesian computation (ABC) [49] as a tool to estimate parameters for dynamical systems describing stem cell differentiation. In cases where conventional likelihood-based approaches fail or are difficult to apply because the likelihood is intractable, ABC methods can provide (approximate) answers.

Most current single cell data provide snapshots of the expression state of a system. Temporal information can typically only be gleaned indirectly, and a host of statistical approaches have been used to remedy this situation [39]. Each of these methods has, however, associated degrees of uncertainty. Here we take a complementary approach where data are assumed to have been generated by a stochastic dynamical system (at different times). We show below that this approach can be used to infer model parameters and determine parameter sensitivities for models of stem cell differentiation. We show that the manner in which we summarise the data – or calculate distances between observed and simulated data – affects the efficiency and reliability of parameter inference profoundly.

## 2 Methods

### 2.1 Waddington Landscapes and Quasi-Potentials for Developmental Processes

Mathematically we can characterize the evolution of cell states, *X*_*t*_, over time *t* by a stochastic differential equation (SDEs),

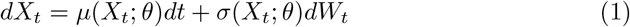

where *µ*(*X*_*t*_; *θ*)*dt* captures the deterministic dynamics of our system parameterized by *θ* and *σ*(*X*_*t*_; *θ*)*dW*_*t*_ depicts the random variations or stochastic properties of this system parameterized by *θ*.

In many cases the focus has been on gradient systems, where we have a potential function, *U* (*X*), such that

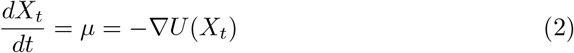

The potential function describes the deterministic dynamics of cell states at each time point and determines the attracting states. The choice of the stochastic dynamics can distort this potential profoundly.

While gradient dynamical systems are flexible and popular – they have also been argued to capture hall-marks of more general developmental systems inspired by developmental biology – they cannot describe all aspects, especially oscillatory and clock-like processes, including the cell cycle [10]. We nevertheless seek to capture many of the dynamics in terms of landscape descriptions, but we make the approximate nature explicit by referring to the mathematical description as the *quasi potential, Ũ*(*X*).

We generally cannot determine *U* (*X*) or *Ũ*(*t*) analytically and instead rely on simulations. We can determine the approximate quasi-potential via the probability density function (p.d.f) *f* (*X*_*t*_), which contains the information about how probable state *X*_*t*_ is. The quasi-potential is then given by *t* [53],

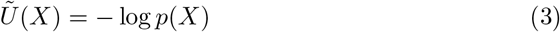

The quasi-potential is thus the logarithm of the probability that a cell is in state *X* [4].

### 2.2 ABC and ABC-SMC

In a Bayesian framework we estimate the posterior probability of a model’s parameters via

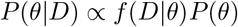

where *f* (*D* |*θ*) is the likelihood of *θ* given the data *D* and where *P* (*θ*) is the prior for the model parameters, *θ*. In ABC the calculation of the likelihood is replaced by a comparison between the observed data and simulated data. ABC methods have the following generic form[2]:

#### Algorithm 1 ABC rejection algorithm

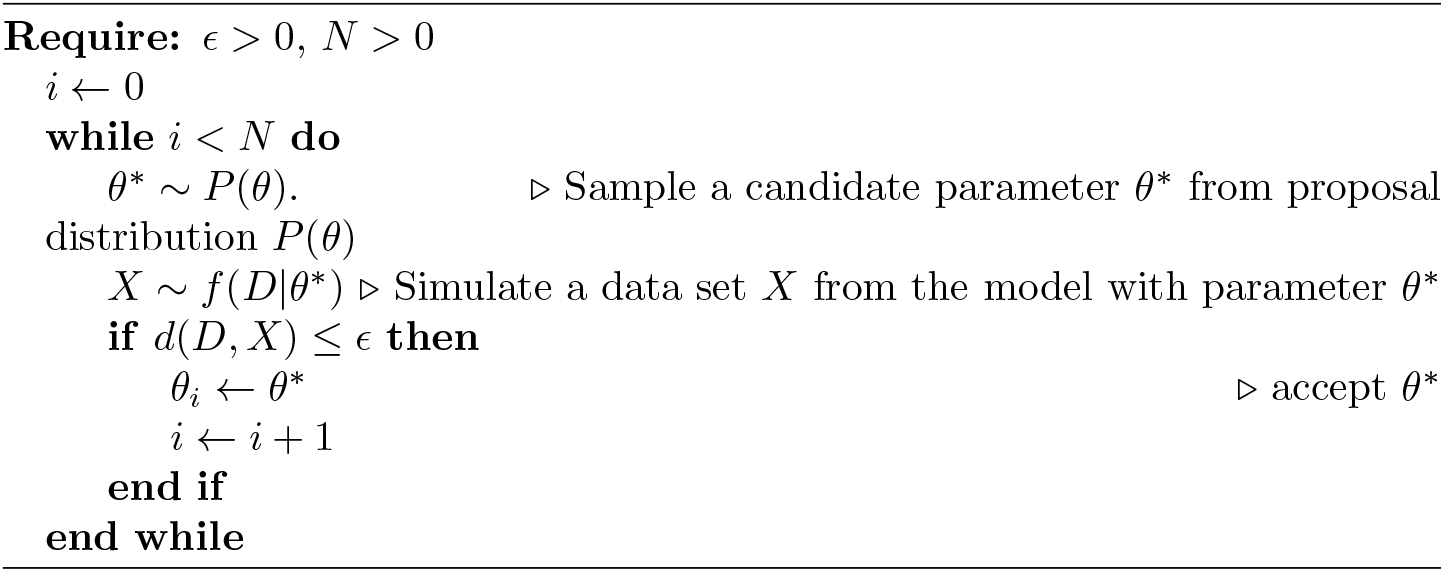

where *d* is a distance function and the tolerance *ϵ≥* 0 defines the intended degree of alignment between *D* and *X*.

The result of any ABC inference is a set of parameter samples from the approximate posterior distribution *P* (*θ* |*d*(*D, X*) *≤ ϵ*). When *ϵ* is sufficiently small then the distribution *P* (*θ*| *d*(*D, X*) *≤ ϵ*) can be regarded as a reliable approximation for the “actual” posterior distribution *P* (*θ*| *D*)[2].

The basic algorithm outlined above, known as ABC rejection, is typically too slow for any problem of interest with more than a few parameter. A range of improvements have been developed in the literature, including Markov chain Monte Carlo, and Sequential Monte Carlo (SMC) approaches. Here we use the SMC approach of Toni et al. [49] adapted to the case of single cell data obtained from by simulations of dynamical systems as the data collected from true developmental systems. In this case a key insight is to find a good distance appropriate for single cell data. We will discuss this in the next section.

### 2.3 Developmental model and simulation procedure

Our analysis will focus on a model that has been used to model embryonic stem cell differentiation processes [15], see Figure 1, where each line represents non-linear inhibition or promotion among these factors. In specific, this model is characterised by the temporal dynamics of four transcription factors: Nanog(N), Oct4-Sox2(O), Fgf4(F) and Gata6(G) where Nanog, Oct4 and Sox2 are complex for maintaining pluripotency and Gata6 is a standard bio-marker for cell differentiation. The detailed representation of their relationship is shown in Figure 1 where each line represents non-linear inhibition or promotion among these factors.

**Figure 1:**
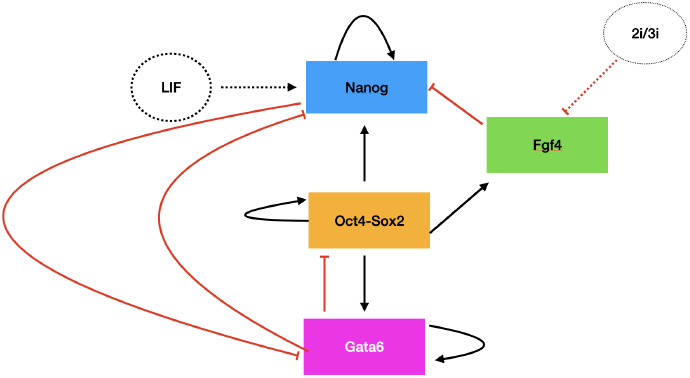
The transcription factor interaction relationship circuit with external factors influences where red line stands for inhibition and black line stands for promotion

We will model the stochastic feature of this processes by implementing a Wiener process on each factor [9].

This system gives rise to two distinct cell states: (i) the pluripotent state is characterised by high values of Nanog and low levels of Gata6; (ii) the differentiated state is characterised by the opposite behaviours for both genes (see Figure 3). The deterministic part of the system is given by [15, 9]

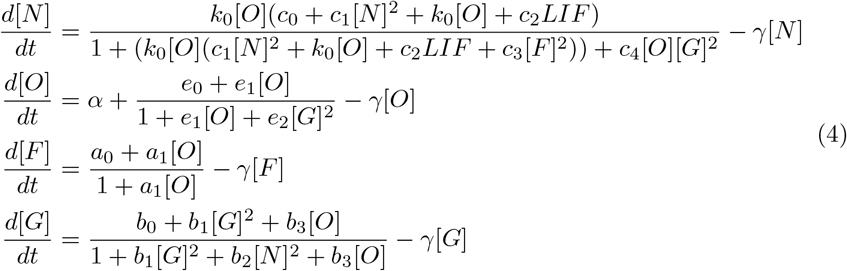

where the value of parameters we will use in this model, according to Chickarmane [15], are:

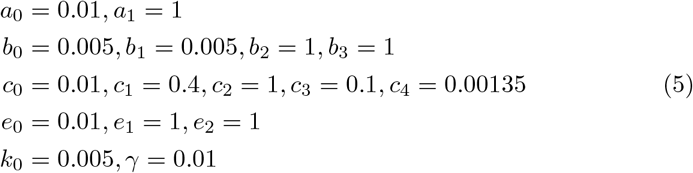

Other than the four transcription factors, there are also some external signals known to influence this system and the propensity functions given above. Leukemia inhibiting factors (LIF) is a cytokine known to inhibit cell differentiation [22] by having a positive effect on the maintenance of Nanog levels. When the level of LIF decreases, the cells will differentiate (Figure 2). Similarly, i2/i3 which are represented by *I*_3_ in our propensity functions, are two different sets of small molecule inhibitors found to be useful in maintain pluripotency in vivo[54]. In our model, increase of I3 will lead to the suppression of Fgf4 which in turn reduces the effect of suppression on Nanog and therefore sustain the pluripotency. Furthermore, this model also includes the possibility of reprogramming by introducing signal *α* in the second propensity function. It can be referred as the reprogramming rate which is in charge of the loop of positive feedback and provides the possible backward transition from the differentiated state back into the pluripotent state: When *α* reaches the critical value, Oct4-Sox2 will induce Nanog and keep raising it towards sufficiently high value and in consequence it will reduce Gata6 level by antagonism[15]. For simplicity, here we only consider the effect of LIF, which is more significant than others, and I3 & *α* are assumed to be zero throughout this analysis. The level of LIF was chosen as fifty for both reference data and simulations in ABC-SMC.

**Figure 2:**
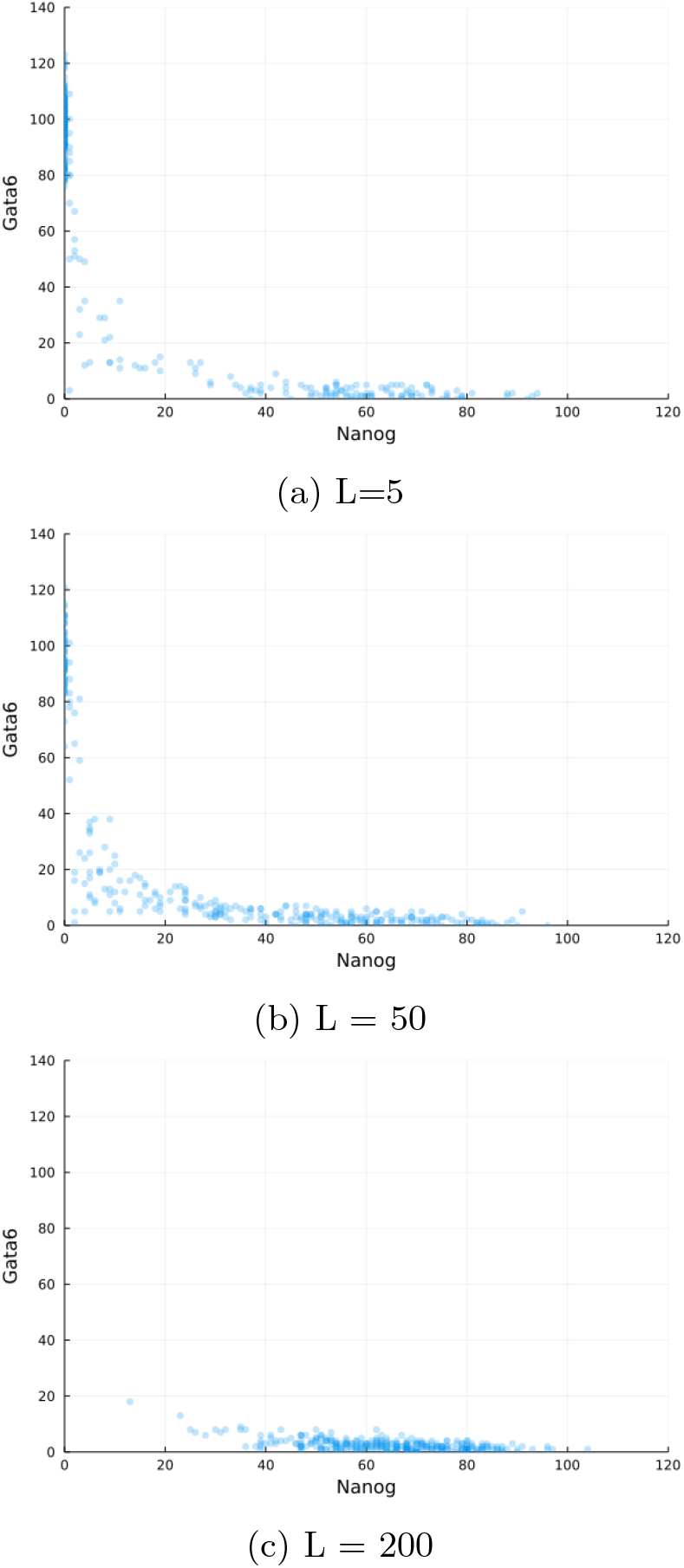
The scatter plots of the relationships between Nanog and Gata6 at the end time for different values of LIF

#### 2.3.1 Simulation procedure

In practice it may be impossible to obtain suitable time course experimental observations. Instead, experimental observations are given as snapshots covering cells at different stages of development or differentiation. To generate our test data we sample cells at ten equally spaced time points. This leads to a dataset that mimics aspects of real experiments [48, 14]. Our ABC-SMC procedure is adapted to this sampling scheme (our reference data and all the simulation data in ABC-SMC were 4 *×*10 *×*300(4 factors; 10 time points and 300 trajectories)). The distance is obtained by summing over all ten time points (i.e. 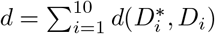 where *D*_*i*_ and 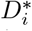are 4 *×* 300 matrices).

For the stochastic simulations (using the Gillespie algorithm[47]) we use the Julia packages *Catalyst*.*jl* and *DifferentialEquations*.*jl*. For the ABC-SMC algorithm we used *ApproxBayes*.*jl*.

We choose the sequence of tolerance threshold *ϵ*_*t*_ based on the 30% quantiles of the population of particles in the last iteration until it reaches the target tolerance threshold *ϵ*_*T*_ or exceeds the set maximum number of simulations (10^6^). That is *ϵ*_*t*_ = *q*_*α*=0.3_(*θ*_*t−*1_), *t ∈* [1, *T*]. We seek to accept 1000 particles of parameters *θ*_*t*_ in each population *t* of the ABC-SMC algorithm. For Kernel Density Estimation with multivariate normal distributions we use *KDEstimation*.*jl* and we set the bandwidth according to Silverman’s rule. And the computation of Optimal Transport is performed in *OptimalTransport*.*jl* in *Julia*.

For all parameters, uniform prior distributions, with lower bound given by 1*/*10 of the true parameters in Equation 5 and upper bounded given by 10 times the true parameters, are used. Since all the data are acquired by simulations rather than from real experimental results, we can choose the final tolerance threshold *ϵ*_*T*_ by simulating data with exactly the same parameters and comparing the distance between them. We run 1000 simulations for the true parameters, 100 with parameters drawn from the priors for each of the distance metrics. All the comparisons are summarized in the density plot, Figure 4. We choose specific *ϵ*_*T*_ values for each distance metric based on these results to correspond to the peak of the density obtained for the true parameters. For example, we choose *ϵ*_*T*_ = 0.46 for the Optimal Transport distance. In practice choice of the *ϵ*_*T*_ schedule is difficult and can affect convergence and speed of convergence considerably [44]; it also depends crucially on the choice of distance metric, and below we discuss a range of suitable distance metrics for single cell data and their relevance for inferring the parameters of models of cell differentiation.

**Figure 3:**
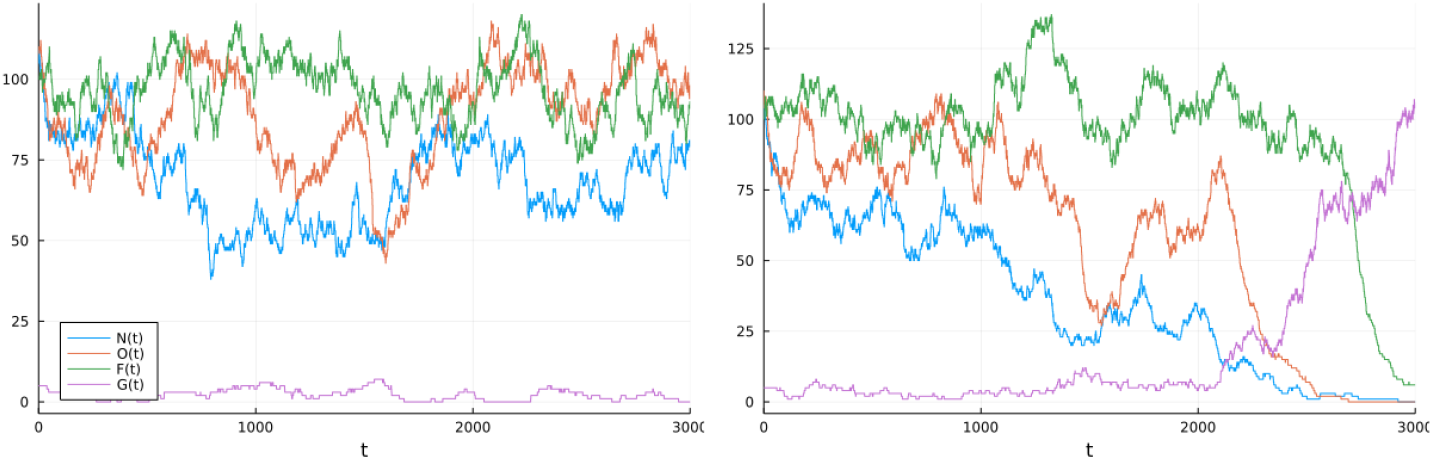
The sample trajectories of the four key transcription factors over time. The left one is Nanog stays pluripotent and the right one is Nanog differentiated into Gata6.

**Figure 4:**
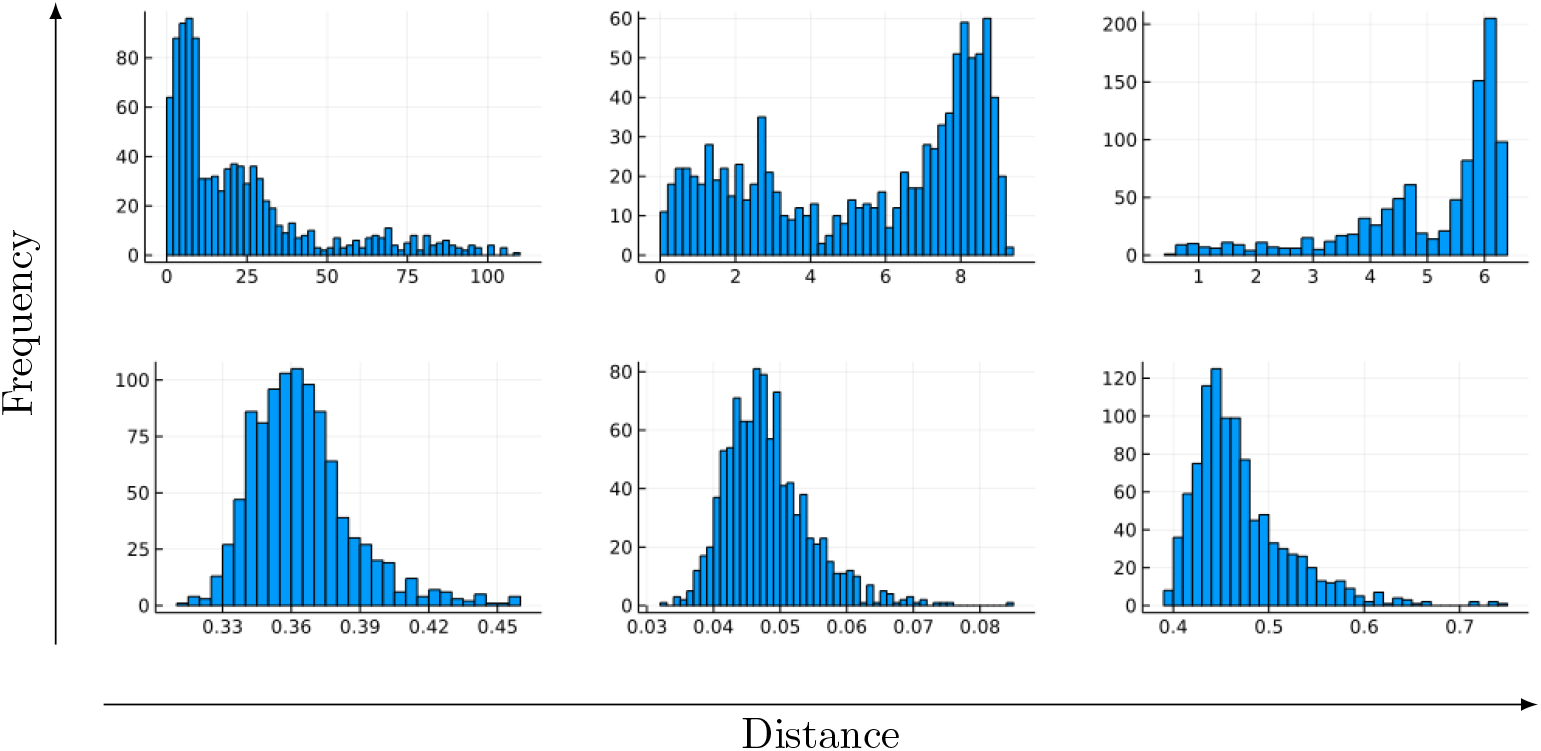
The distribution of distances of 1000 comparisons for different distance metrics. Top first: Bhattacharyya distances with parameters drawn from the priors. Top second: MMD with parameters drawn from the priors. Top third: Optimal Transport with parameters drawn from the priors. Bottom first: Bhattacharyya distances with identical parameters. Bottom second: MMD with identical parameters. Bottom third: Optimal Transport with identical parameters.

## 3 Results

### 3.1 Distances for Single Cell Data

In Eqn. (3) we determine the quasi-potential *Ũ*(*X*) from the probability distribution over cell states, *X*. For our ABC SMC procedure we therefore need suitable distances between probability distributions. In this section we examine and compare some measures that quantify the discrepencies between probability distributions, as well as their underlying geometric properties before adopting them in our ABC-SMC inference scheme.

#### 3.1.1 KL-Divergence

We start with the KL-Divergence which is one of the simplest and most commonly used measure in comparing probability distributions. For data *D ∼ Q* and model distribution *P*_*θ*_, the KL-divergence is defined as,

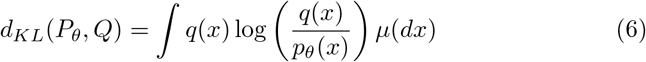

Here *q* and *p*_*θ*_ are the corresponding probability density functions with respect to the same probability measure *µ*. The KL-divergence is closely related to maximum likelihood theory [51]:

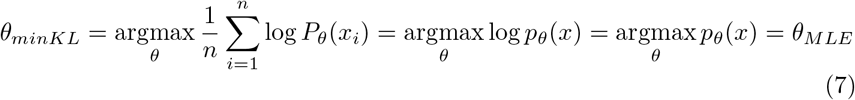

However, if there exist *x*_*i*_ such that *q*(*x*_*i*_) = 0, the KL-Divergence diverges to infinity which can become problematic in numerical computations. Adding a small amount of random noise may help, but at the risk of adding errors. Because of the numerical issues we prefer other measures of the distances between probability distribution.

#### 3.1.2 Bhattacharya Distance

The Bhattacharyya distance is used in signal selection and pattern recognition[41] and defined by,

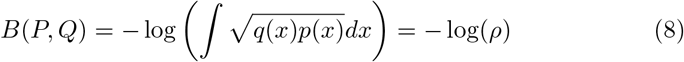

for distributions *P, Q* with probability density functions *p*(*x*) and 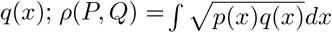 is also known as the Bhattacharyya coefficient. Unlike the KL divergence, the Bhattacharyya distance is symmetric and unbounded (0 *≤ B≤ ∞*).

An advantage of the Bhattacharya distance is that it has been proved that for any two sets of parameters *θ*_1_, *θ*_2_ if we have *B*(*θ*_1_) *> B*(*θ*_2_) then there exist *π* = (*π*_1_, *π*_2_) prior distributions that satisfy *P*_*e*_(*θ*_1_, *π*) *< P*_*e*_(*θ*_2_, *π*) where *P*_*e*_(*θ, π*) is the error probability with parameter set *θ* and prior probability *π*[26]. This property makes it particularly meaningful for parameter estimation in an ABC framework while considering the optimal Bayes error probabilities. Furthermore, this metric gives good results for Gaussian noise problems, which is useful for SDE problems modelled that use standard Brownian noise terms [41].

Even though this criterion works for most common distribution functions, there are two drawbacks in our setting,

1. Our model is implicit (i.e. the data points are generated by a set of equations describing the biophysical processes), but the corresponding distributions cannot be represented by a density function without using kernel density estimation (KDE) or a different density estimation methods in high-dimensional space. However, due to the curse of dimensionality, the number of data points required to have accurate estimates increases dramatically with increasing dimensionality[33], distance metrics which require KDE would be problematic not only in terms of accuracy but also efficiency.
2. This metric only measures the difference of probability density in a point-wise way which ignores the underlying geometry name of the probability space[21].

Therefore, we next introduce distance metrics that do not require KDE and which take the geometry of feature space into account.

#### 3.1.3 Optimal Transport

Next we consider an optimal transport distance metric that is free of density estimation by comparing samples directly and maintains the underlying geometry in feature space. This metric has been widely used in the field of machine learning and image recognition for its efficiency [17]. Intuitively, the key to Optimal Transport is to figure out the minimal cost that is required to transform one image into another, or in our case, one probability distribution into another.

Suppose we have two probability distributions *P, Q* on the probability space*χ*, Γ with probability measure *µ, ν* respectively. Let *T*_#_*µ* denote the forward of *µ* by *T*. That is

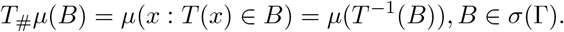

The original version of optimal transport is defined by [45]

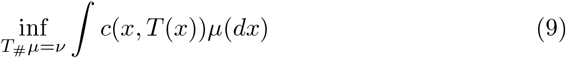

where *c* : *χ×* Γ↦ [0, *∞*) is a measurable function on *χ×* Γ known as the cost function.

However, this technique is clearly not applicable when there does not exist any forward equation *T* such that *T*_#_*µ* = *ν*. To overcome this, we adopt Kantorovich’s version of Optimal Transport or the *p*-Wasserstein distance *W*_*p*_(*P, Q*), defined as [45],

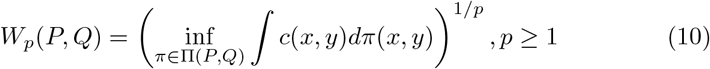

where Π(*P, Q*) represents all the joint probability measures on *χ ×*Γ with marginal measures, *µ* and *ν*, respectively. This version is known to be well defined in general scenarios[45].

If in addition we have *f*_*P*_ : *χ ↦*ℝ and *f*_*Q*_ : Γ↦ ℝ bounded and measurable functions with

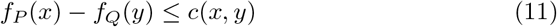

it is easy formulate the duality form of Optimal Transport [1]

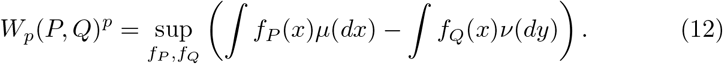

Since we are working on the same probability space for both *P* and *Q*, say *χ* = Γ, we can have the generalized form under 1-Lipschitz (Lip1) continuous functions on *χ* [8],

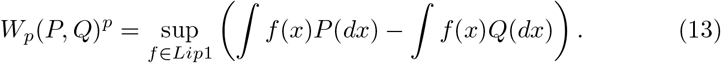

If we have two different probability distributions, say *P, Q ∈Pχ*, then the mixture distribution

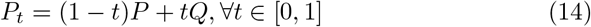

can be considered as a curve connecting *P, Q* under the whole space of probability distributions, *P*_*χ*_. It can be shown that for *p* = 2, the Wasserstein distance will give a minimal geodesic under such a probabilistic space of distributions[8]. In other words, whenever Optimal Transport proposes a probability distribution that transports from *P* to *Q* under *χ*, we will have the shortest path or distributions between *P* and *Q*. This result gives an extremely meaningful insight in our ABC-SMC framework to compare samples and simulations in geometry. It is worth noting that there may exist many minimal geodesics given by the Optimal transport which satisfy Equation (14).

#### 3.1.4 Maximum Mean Discrepancy

We also consider the Maximum Mean Discrepancy (MMD) measure which has been used for of goodness of fit tests [27], and in an ABC framework [3]. The principle here is to find a kernel function that will give different expected values from two samples when they are not drawn from the same distribution. The underlying rationale is that the evaluations of such function at sample points, which are drawn from different probability distributions, can provide enough information about their difference[7].

The initial definition of MMD is as follows: Let ℱ be a class of functions *f* : *χ↦* ℝ and *P, Q* be distributions of samples from *X, Y* which are independent and identically distributed with respect to *P* and *Q*; then MMD can be defined as

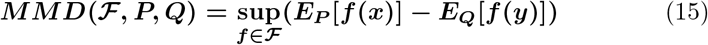

It is clear that if and only if *X, Y* are drawn from the same distribution (i.e. *P* = *Q*) then the MMD is zero for any functions *f∈*ℱ ; but if the class of functions ℱ is too “large”, the statistic would be much greater than 0 for most of the finite samples *X, Y*, which “strengthen” the differences in distribution.

In order to avoid this scenario while simultaneously allowing reasonable discrepancies between *P*&*Q* to be detected, there must be a suitable restriction on the class of function ℱ. The trade-off is achieved by restricting ℱ to be the unit ball in reproducing kernel Hilbert space (RKHS)[7]. The MMD can be calculated from this space by calculating the expected values of this kernel function under distributions *P, Q*. Let ℋ be a RKHS on feature space *χ* induced by a feature map *ϕ* : *χ ↦*ℝ. The kernel is given by the inner product between the feature maps under RKHS,

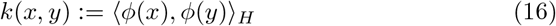

It is guaranteed that MMD can detect the difference between distributions *P, Q* of samples from *X, Y*. When ℱ restricted as the unit ball in the reproducing kernel Hilbert space (i.e. ℱ := *f* : *χ↦*ℝ ||| *f*||_ℋ_ *≤* 1), then *MMD*(ℱ, *P, Q*) = 0 if and only if *P* = *Q*[7]. With characteristic kernels (i.e. kernel mapping is injective), MMD is proven to be a probabilistic metric which takes values in distribution functions [46]. A standard choice of this is the Gaussian kernel,

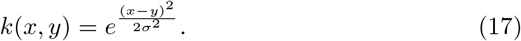

We will use the median heuristic to set the bandwidth of Gaussian kernels with 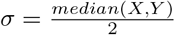.

In practice, given samples from *X* with sample size *m*, and from *Y* with sample size *n*, it is straightforward to determine the corresponding MMD,

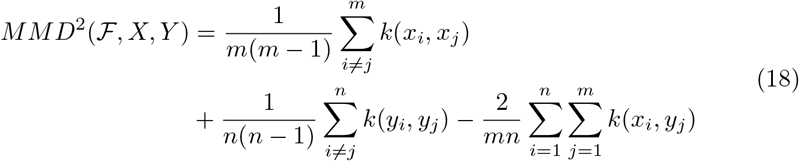

Since MMD can be easily computed by empirical observations, adopting it with ABC-SMC simulators would be very efficient.

MMD is defined similarly to Optimal Transport 13 under different sets of functions ℱ. Such metric problems are known as Integral Probability Metrics(IPMs)[32]. It has been shown that all such IPMs preserve the minimal geodesic property and it has further been proved that there exists a unique minimal geodesic under MMD in comparison to Optimal Transport (For details see Theorem 5.3 in [8])

### 3.2 ABC inference depends on distances

The obtained approximate posterior distribution of all the parameters for each distance metric are summarized in Figure 8,9,10. Our results of posterior distributions suggest a good agreement in estimating parameters using ABC-SMC with different metrics. With regard to the identifiability of parameters, we figure out that there is a well structured shape of probability distributions for most of the parameters (*c*_1_, *c*_2_, *c*_3_, *e*_1_, *b*_1_, *b*_2_, *b*_3_) but other parameters are more likely to have “flat” distributions which are not well inferable.

Here we test the accuracy of the estimated posterior distributions by drawing particles from such posterior distributions and see if we can reproduce the dynamic trajectories of our system. The simulation result using drawn particles from ABC-SMC results with different distance metrics is given in Figure 5 and it is much akin the pattern of dynamics as in reference data (Figure 2b). Our result suggest both, the applicability of ABC-SMC in parameter estimations from single cell data, and the importance of choosing an adequate distance measure.

**Figure 5:**
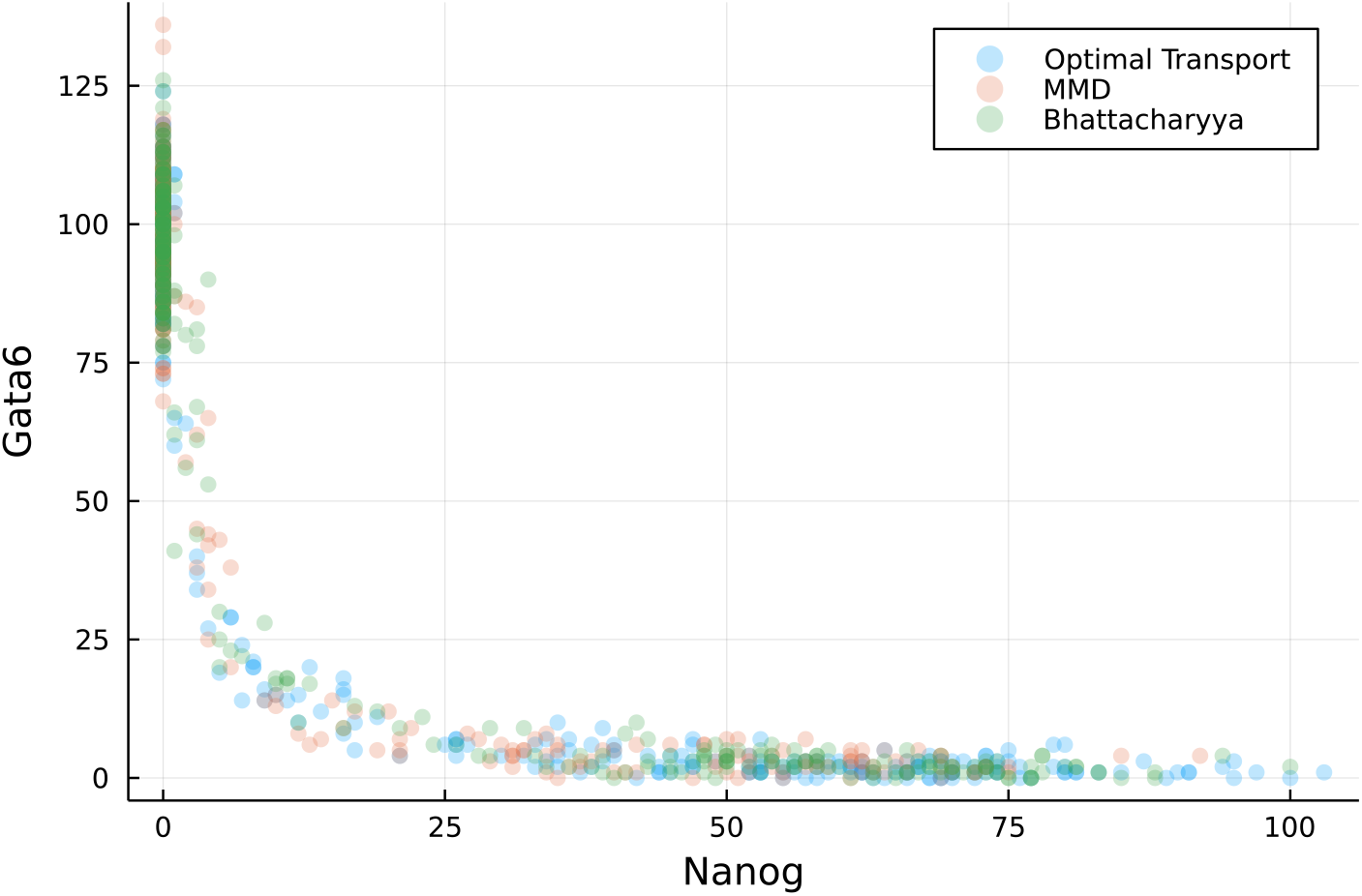
The plot of Nanog versus Gata6 at final time point using sampled particles from the last population; LIF was set as 50

We choose the sequence of tolerance thresholds for MMD as the 30% percentile, which here gives rise to a schedule of,

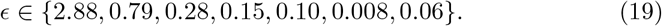

which decreases slowly, see Figure 4. For comparison, the sequence of tolerance thresholds for Optimal Transport is

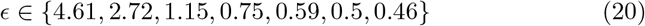

which decreases constantly even after reaching the the boundary support of density for true parameters in Figure 4. Our result indicate that it in an ABC-SMC context it might be easier for Optimal Transport to take an efficient path to parameter inference than MMD. Theoretical reason for that would be worth studying in the future.

The ABC-SMC posterior distribution helps to analyze the relationships among all the parameters in our system. We can determine underlying correlations between pairs of parameters in the pair-wise plots (Figure 8). We find that *c*_1_ and *c*_3_ are strongly correlated with a clear linear trend and some slight linear relationships are also shown between *e*_1_ and *e*_2_ and *b*_1_ and *b*_2_. With regards to the identifiability of relationship between *b*_1_ and *b*_2_, we find that the performance of the Bhattacharya distance lags behind those of other distances (Figure 10).

Many such dynamical, including biological models, are know to exhibit “sloppy” behaviour: the dynamical behaviors of such systems are usually controlled by a small number of parameters [50, 19]. In addition to that, varying correlated parameters will still reproduce the same exhibiting behaviors of such systems. For example, in our result, it would be easy to determine the value of *c*_3_ or *c*_1_ if the other one is known while reproducing the same trajectories of transcription factors due to the correlation between them. We will show that our approach can help resolve this problem by identifying such set of correlated parameters from joint distribution and the detail of this “sloppy” analysis will be discussed in the following section.

With regard to computational cost, we found that MMD is the fastest one and Optimal Transport follows as reported in the theoretical time complexity[35]. And with Bhattacharya distance, it takes more than three times amount of time as in MMD where most of them are consumed in probability density estimation by KDE. Therefore, our results suggest the priority of density free methods in our ABC-SMC framework in single cell model.

### 3.3 Parameter Sensitivity

Visualizing and analyzing the sensitivity of parameters that are in high dimensions is typically hard and so does in our joint posterior distributions. To counterpart this, the most simplest and convenient way is directly focusing on the variance of posterior distributions and so forth adopting Principle Component Analysis (PCA) to measure the variability of parameters [5, 42].

Starting with the posterior distributions of our parameters obtained from ABC-SMC, we can use the traditional Method of Moments to estimate the sample covariance matrix Σ:

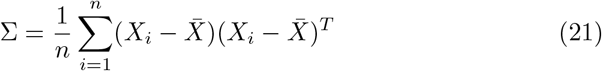

And we can therefore estimate the variance of each parameters after applying the spectral decomposition to Σ:

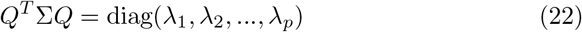

where *Q* = (*q*_1_, …, *q*_*p*_) is the eigen-parameter (vector). This eigenvector forms a orthogonal matrix and *λ*_1_, *λ*_2_, …, *λ*_*p*_ are the corresponding variances of each parameter which are known as the principal components of Σ. Therefore we can easily measure the variability of data explained by each parameter by diagonalizing the covariance matrix and perform the sensitivity analysis.

If *q*_*j,k*_ is the element of direction associated to the *k*^*th*^ parameters then we will have the projection of *θ*_*k*_ through the inner product,

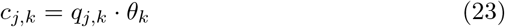

where *c* = (*c*_1_, …, *c*_*p*_) are such principal components. We then have

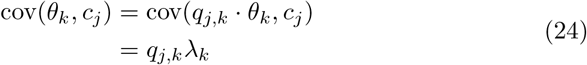

which gives the correlations between principal components *c*_*j*_ and eigen-parameters *θ*_*k*_. Thus, we can use the standardized square correlations:

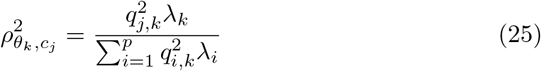

to interpret the proportion of variance eigen-parameter *θ*_*k*_ explained by principal component *c*_*j*_. From this we can use the total proportion of variability explained by parameter *θ*_*k*_ to quantify the sensitivity of parameter *θ*_*k*_ by,

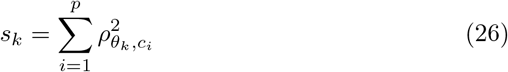

We note that using PCA to analyze sensitivity is the same as eigenvalue analysis of the Hessian matrix around the MLE or the equation of least square errors [42].

We can define the log posterior density by

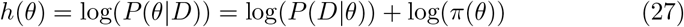

And using Taylor expansion of logarithm along with the log posterior distributions, we obtain [42],

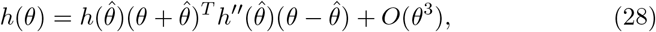

where 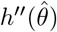 is the Hessian matrix of log posterior distribution as in Equation **??**.

When assuming the parameters are drawn from the multivariate normal distribution we can easily deduce the asymptotic covariance matrix of *θ* by the Fisher information matrix:

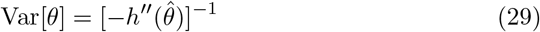

From this, we can conclude that the eigenvalues of the Variance-covariance matrix *λ*_*k*_ in Equation 22 and the eigenvalues of the corresponding Hessian matrix *v*_*k*_, which results in asymptotic variance by assuming Multivariate Normality, are inversely related[28]:

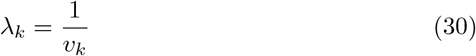

which legitimates the statistical property of our PCA result.

We summarize the result of variability/sensitivity of each parameter in Figure 6 following by the steps above. It is known that for change of value in sloppy parameters will alter the system behaviors slightly and change of that in stiff parameters will lead to dramatic change in dynamics. In terms of paramerter variance, if we characterize parameters that explain more than 1% of the variance for the posteriors as sloppy and otherwise as stiff we can conclude that parameters *a*1, *b*2, *b*3, *c*1, *c*2, *e*1, *e*2 are the sloppy parameters with higher uncertainties in our system.

**Figure 6:**
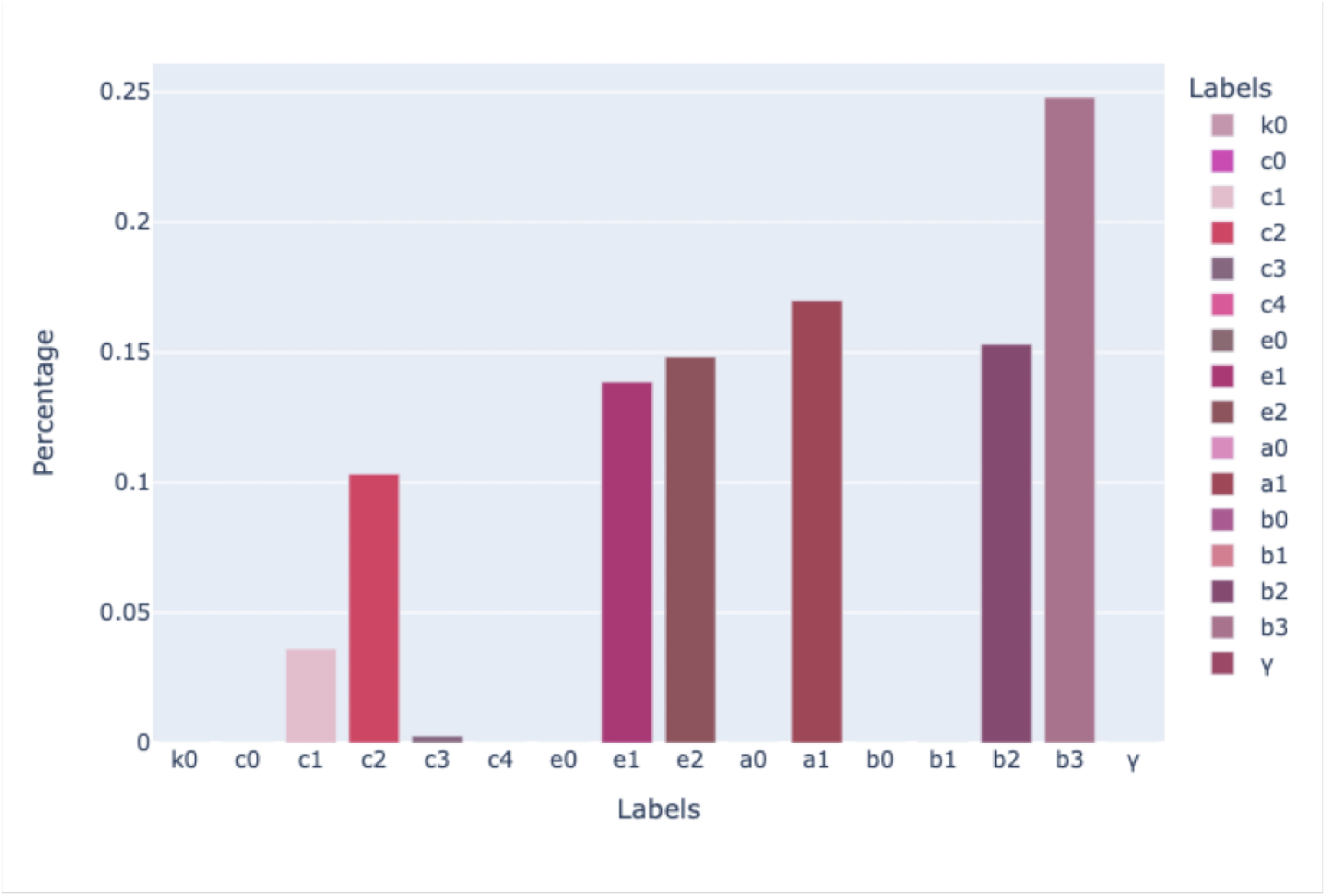
Variability of the model explained by different parameters

Our result also suggests that all the regeneration rates(*c*_0_, *e*_0_, *a*_0_, *b*_0_) as well as the constant degradation rate *γ* are more stiff than others. This is not quite surprising since these parameters actively decide the extent of change in each factor throughout the whole time. For gene transcription factors Oct4-Sox2 and Fgf4, all other parameters are sloppy except their corresponding regeneration rates and the constant degradation rate *γ*. This result has an agreement with that higher variance in Oct4-Sox2 and Fgf4 along with the time are expected in contrast to the others as shown in [10]. In view of the scaling parameters *k*_0_, *c*_4_ for Oct4-Sox2 and Gata6 which are stiff in factor Nanog, we found that the effect of Oct4-Sox2 and Gata6 on Nanog and hence on the fate decision making in our system is very distinct. In contract, the effect of Nanog on Gata6 is less based on the sloppy scaling parameter *b*_2_. As a compensate, self-regeneration or degradation rate *b*_1_ is much more significant on Gata6 which justifies its role as bio-marker of differentiation.

## 4 Discussion

The inference of parameters from experimental single cell data for complex systems is of significant practical interest. While there are many previous approaches based upon different Bayesian framework or ABC algorithms[38, 13, 6, 38, 11, 28], summary statistics coupled with some simple distance metrics like the Euclidean distance of *L*^*p*^ norm are commonly chosen[18]. However, recent studies, inspired by Waddington’s landscape, has revealed that the existence of potential provides a meaninful description of the system;s dynamics and is quantitatively associated with the probability distribution over states *X*_*i*_ [53, 4, 9]. Therefore, we argue that employing such basic distance metrics will result in information loss due to non-sufficient summary statistics.

In this study, we investigate parameter inference through the ABC framework using different distance metrics for a specific biological system. We explore the effectiveness of three metrics: Maximum Mean Discrepancy (MMD), Bhattacharya distance, and Optimal Transport. Our results have shown that the inference of parameters based on these metrics can successfully capture the dynamics of this biological system.

To see the significance of the quasi-potential landscape as a significant factor in determining model parameters within the ABC paradigm, we conducted a comparison by employing our ABC-SMC algorithm with the simple Euclidean distance metric. Specifically, we considered two types of summary statistics: the first and second moments (*E*(*X*), *E*(*X*^2^)) and the logarithm-transformed probability of cells in pluripotent states 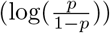 where *p* was determined using K-means with 2 classes.

We summarized the results of the posterior predictive distribution check using 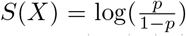 as the test statistic (Figure 7). We outlined the extreme quantity *p* (*Pr*(*S*(*X*) *> S*(*D*))) and Mean Squared value (MSE) and observed that the posterior distributions obtained using distance metrics that compare probability distributions were more consistent with the simulated experimental data, outperforming those obtained with the Euclidean distance. Therefore, we concluded that accessing information from the landscape is crucial for determining the parameters of dynamical systems. To achieve this, it is necessary to adopt distance metrics capable of quantifying the differences between intrinsic probability distributions. The result of this comparison highlights the importance of considering the landscape structure and employing suitable distance metrics for accurate parameter inference within the ABC framework.

**Figure 7:**
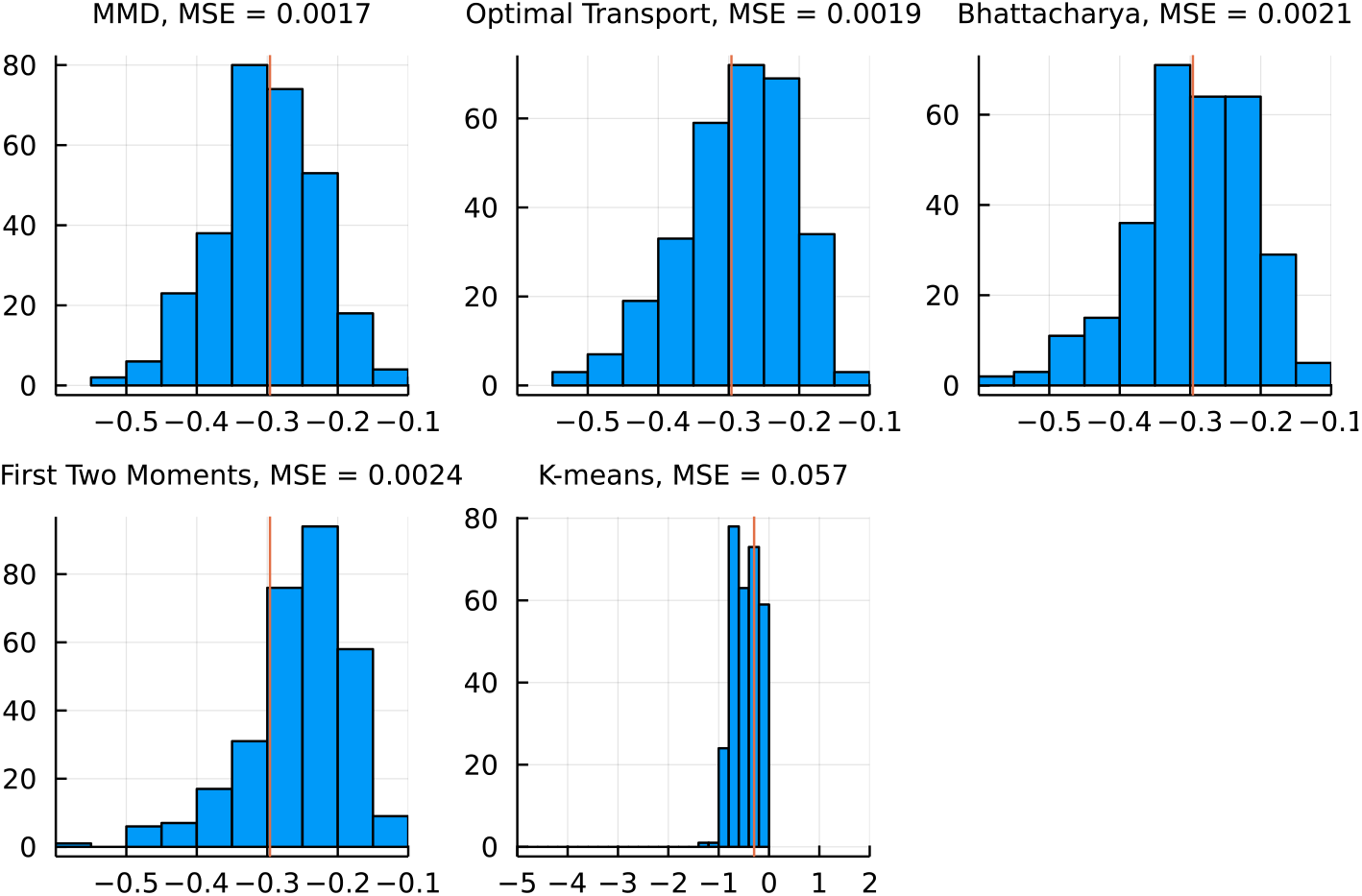
Realized vs. posterior predictive distributions for ABC-SMC results obtained with the use of different distance metrics: 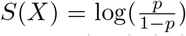 (where *p* is the probability stem cell ends in pluripotent state), compared to simulations from the posterior predictive distribution for each distance metric. The vertical line presents the test statistics from simulation of true parameters and the corresponding Mean Squared Error (MSE) for different metrics are presented.

Our findings suggest that among the evaluated metrics for comparing distributions, MMD yields the highest accuracy. It is known that Optimal Transport and MMD belong to the same category of metrics known as Integral Probability Metrics (IPMs) [32]. Moreover, when they both use the same kernel functions, Optimal Transport can be considered as an interpolation of MMD [35]. Common knowledge suggest that Optimal Transport would perform better in determining the target estimate, especially when employing gradient flows such as in the gradient descent approach [21]. However, our results show a different outcome, with MMD exhibiting better performance in our framework. Therefore, our result suggest that MMD is more advantageous for applications in non-gradient approaches, like in the ABC (Approximate Bayesian Computation) scheme.

With regard to computational cost, both MMD and Optimal Transport metrics are quite similar, and they outperform Bhattacharya distance significantly. The key reason for the computational difference with Bhattacharya distance lies in the estimation of the probability density function through the KDE method. This drawback would become more significant as the dimensionality increases. Considering the computational difference, we strongly recommend using MMD as the distance metric where samples are directly compared without density estimation and implicitly quantifying the differences between distributions. This property makes MMD computationally efficient and well-suited for dealing with high-dimensional data and it offers a more practical and efficient option for parameter inference with ABC algorithm in dynamical systems.

Although the potential landscape offers valuable insights into the dynamics of biological systems, it is not the sole factor governing these processes. Conventionally, the quasi-potential is defined by taking the negative logarithm of probability distributions, following Boltzmann’s rule, assuming a static and time-homogeneous landscape [34]. However, recent research has delved deeper into this field, revealing that cell systems are more like time-inhomogeneous, and highlighting the existence of transient landscapes [20, 9].

To accurately capture the time evolution of stem cells, it is essential to consider the curl of probability flux, which interacts with the potential landscape to govern the ongoing trajectories of cell fates [29]. In our approach, although we attempted to quantify the variational structure of the landscape by collecting data points at different times, we still lost the information related to the probability flux.

Since we have demonstrated the significance of landscape information in inferring parameters from single-cell data, we believe that our framework can be further enhanced and better understood by incorporating and measuring the probability flux under appropriate distance metrics. This extension would allow us to gain a more comprehensive understanding of the underlying dynamics and improve the accuracy of parameter inference in complex systems.

## Acknowledgment

YL and MPHS gratefully acknowledge funding from the ARC through an ARC Laureate Fellowship to MPHS (FL220100005); SYZ acknowledges funding from the Australian Government Research Training Program and an Elizabeth and Vernon Puzey Scholarship. ITK was funded through a Wellcome Trust PhD studentship.

## 5 Appendix

**Figure 8:**
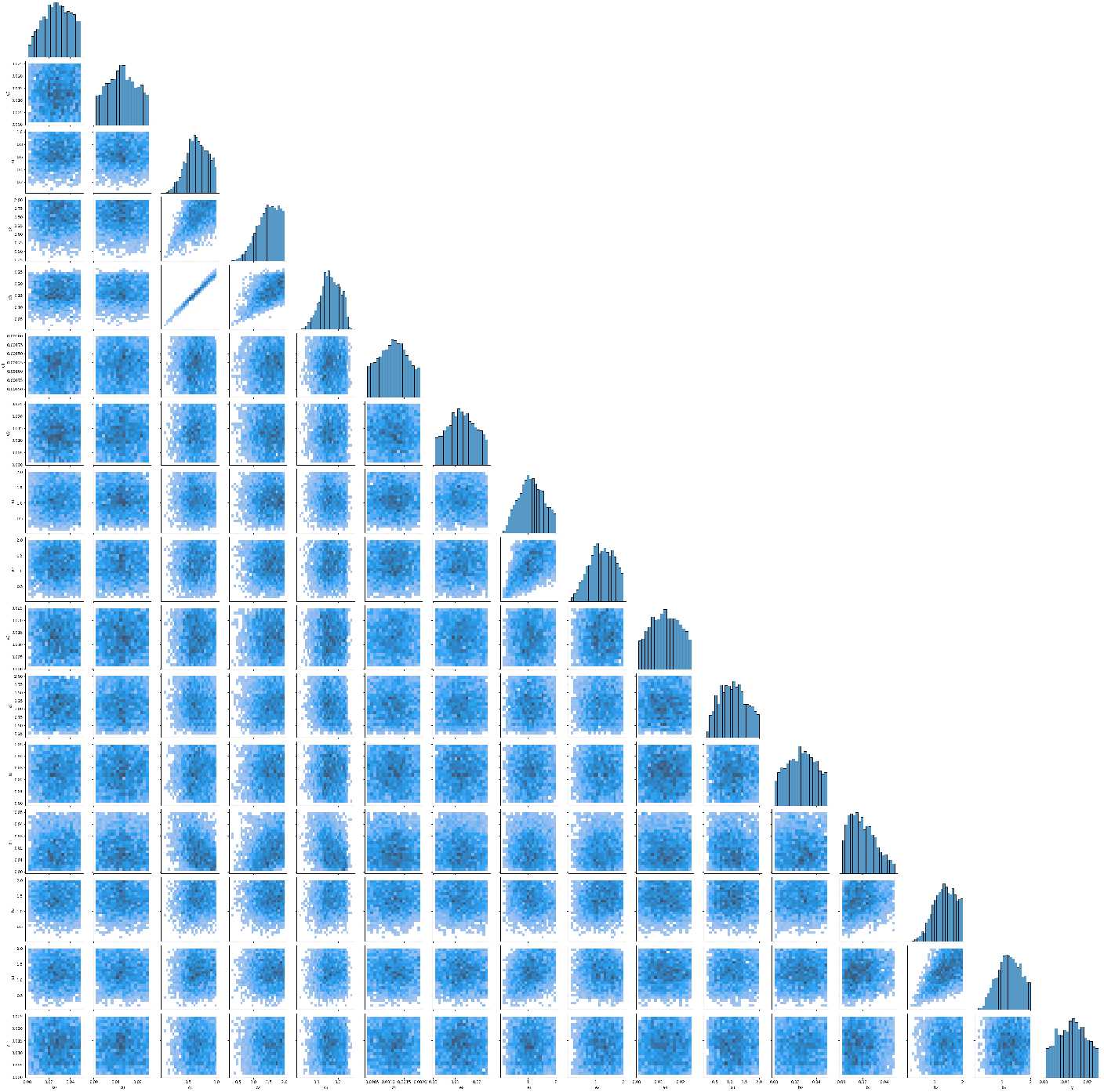
Results of ABC-SMC in parameter estimations for the stem cell model using Optimal Transport. Diagonal: Histogram of each parameter. Lower triangle: 2D pairwise scatter plot of the joint posterior distributions between two parameters.

**Figure 9:**
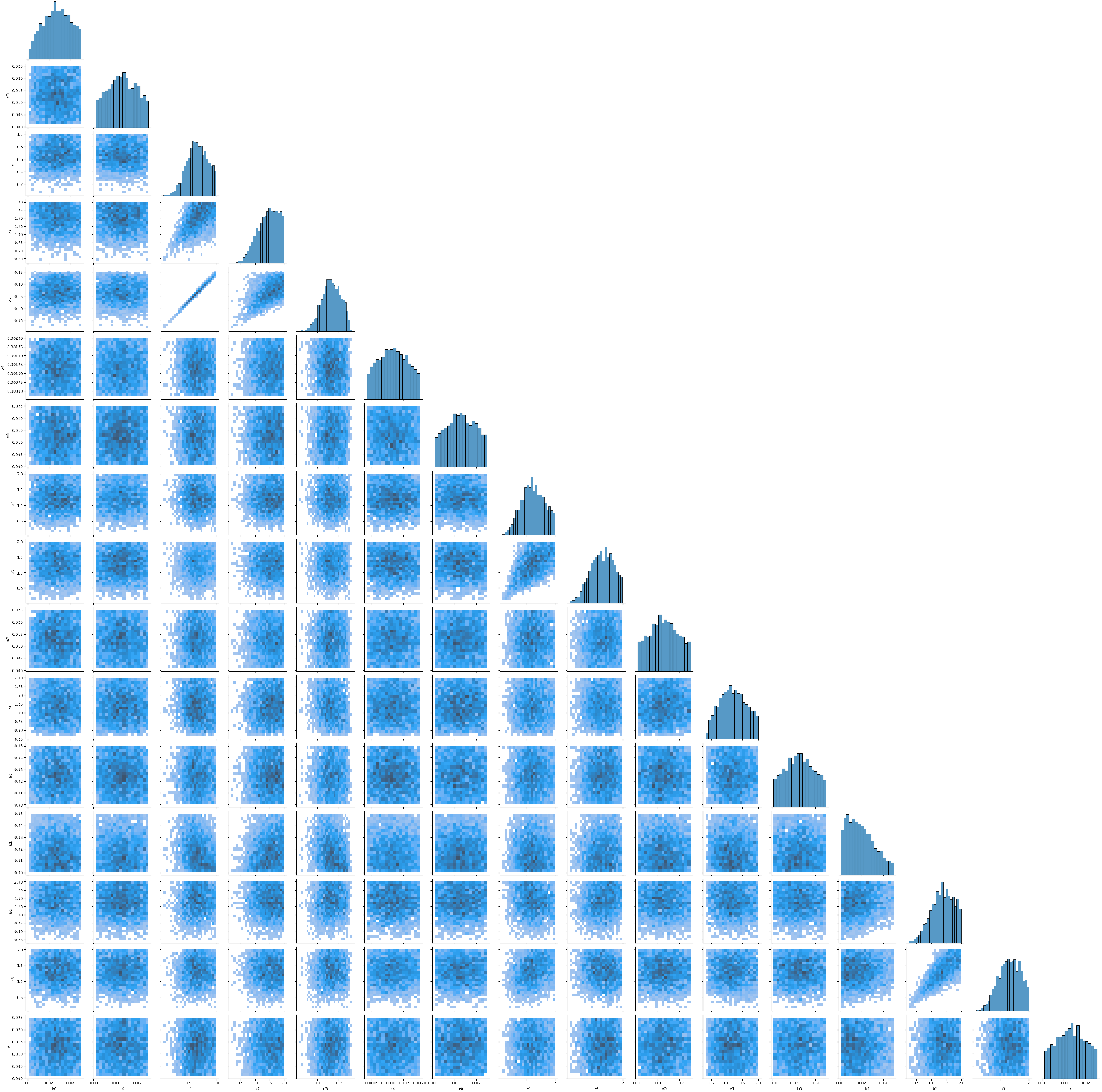
Results of ABC-SMC in parameter estimations for the stem cell model using MMD, similar as Figure 8

**Figure 10:**
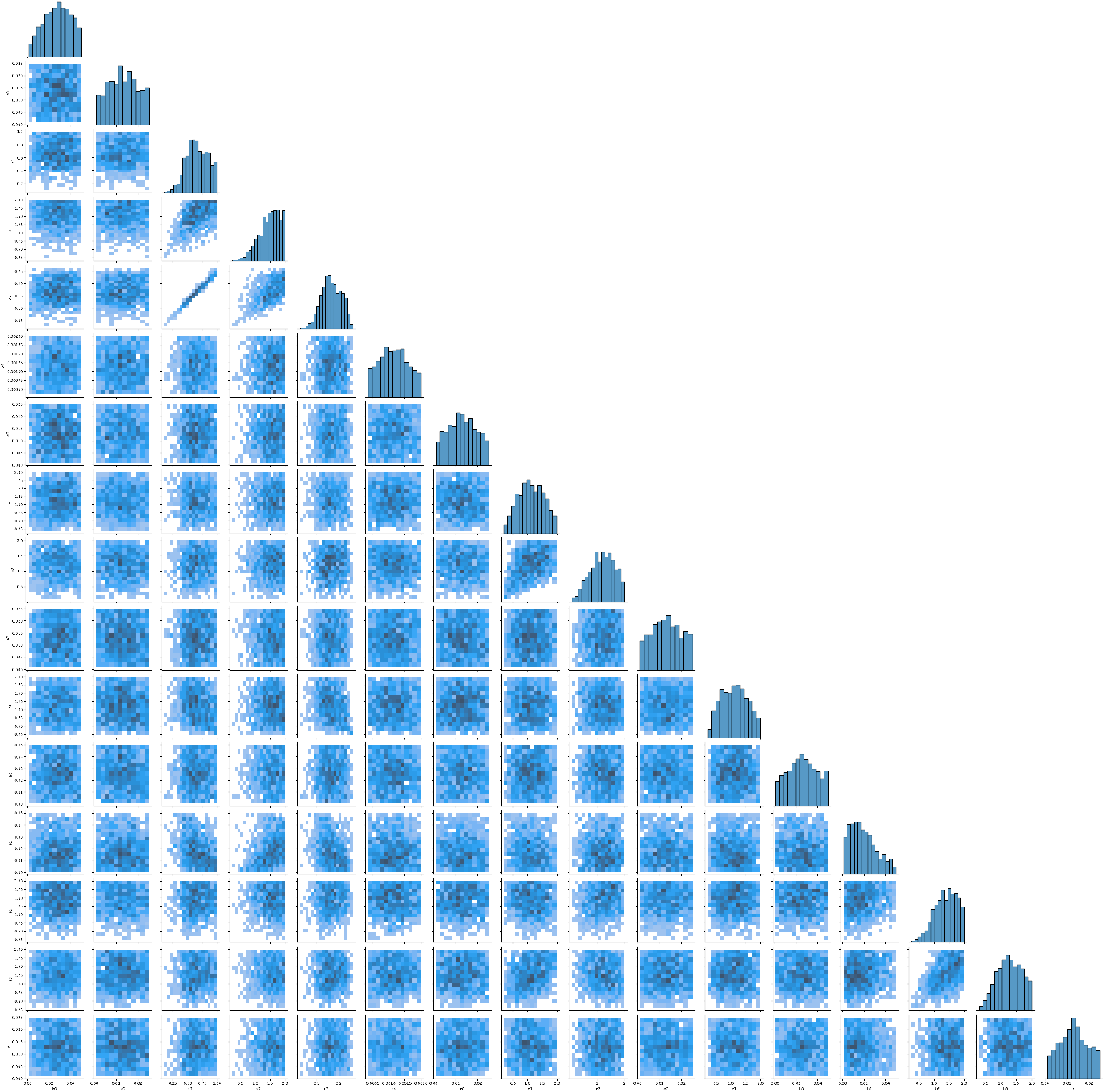
Results of ABC-SMC in parameter estimations for the stem cell model using Bhattacharya distance, similar as Figure 8

## Notes

### Competing Interest Statement

The authors have declared no competing interest.

